# A Ponceau Staining based Dot-Blot Assay for reliable and cost-effective Protein Quantification

**DOI:** 10.1101/764712

**Authors:** Dario-Lucas Helbing, Leopold Böhm, Yan Cui, Leonie Karoline Stabenow, Helen Morrison

**Affiliations:** Leibniz Institute on Aging, Fritz Lipmann Institute, 07745 Jena, Germany; Faculty of Medicine, Friedrich-Schiller-University Jena, 07743 Jena, Germany; Institute of Molecular Cell Biology, University Hospital Jena, Friedrich-Schiller-University Jena, 07745 Jena, Germany

**Keywords:** Dot-Blot, Ponceau S, protein quantification, cost-effectiveness, sustainability

## Abstract

Reliable quantification of protein extracts from tissues can be a challenge e.g. due to interference of the high fat content in tissues of the nervous system. Further problems like under- or overerstimation of protein concentrations in protein quantification kits like the bicinchoninic acid (BCA) assay can occur. In addition, common lysis buffers such as RIPA buffer are known to be unable to solubilize a large amount of proteins (~10-30%) leading to unsatisfactory and unreliable experimental results with techniques such as immunoblotting. In this work, we have developed a Ponceau S staining based protein quantification assay. This assay is compatible with tissues or cells directly lysed in 2x SDS gel loading buffer, containing bromophenolblue, leading to more complete protein extraction. Protein concentrations of several samples can be determined in a fast and cost-effective manner and subsequent experiments (e.g. Western blot) can be performed without loss of proteins. The presented protein quantification method is highly reliable, fast and economical. Using this method, it is possible to save between 2300 to 3200€ per 1000 lysates as compared to the costs of a commercial BCA kit.

## 1. Introduction

A variety of different methods exist for quantification or estimation of total protein content in lysates from cells and tissues. However, the most common methods like the BCA(1), Lowry(2) or Bradford(3) assay are based on photospectrometry which has the disadvantage of fast saturation problem if the protein concentration in lysates is high and therefore outside the range of the standard. Additional problems may occur because a variety of chemical substances (e.g. SDS, although SDS-containing buffers are used for tissue lysis and subsequent quantification with BCA assay) commonly used for effective lysis of biological material and consequent solubilization of the extracted proteins are known to interfere at high concentrations with the chemical reactions underlying the aforementioned methods(1, 4). Furthermore, high concentrations of lipids (e.g. in nerve and brain lysates) are known to interfere with photocolorimetric assays like the BCA assay(5). In addition, large volumes of lysate may be needed leading to a profound loss of the sample.This is an undesirable side-effect especially if handling small tissues with small protein yield like sciatic nerves. Thus, these methods are not only costly, likely unreliable and require large sample volumes. Here, we demonstrate a (compared to the commercial BCA kit) time- and money-saving method for protein quantifications with the use of small sample volumes. We achieved this by using a Ponceau S based Dot blot method (“PDB-assay”). Ponceau S staining is normally used as a loading control for protein loaded membranes during Western blotting(6). Our assay gives completely linear standard curves and shows no saturation even in the range of very high protein concentrations of BSA (around 8 μg/μl) and also when testing samples with high protein content (e.g. spleen, brain).

## 2. Materials and Methods

### 2.1 Reagents

Ponceau S (Merck KGaA Darmstadt, Germany, #P3504-10G)

Pierce Bovine Serum Albumin Standard (BSA) ampules Thermo Fisher Scientific Inc., Waltham, MA, USA, #23209)

2x SDS Gel Loading buffer (=”2x SDS LB”):

100mM Tris-HCl (Carl Roth GmbH + Co. KG, Karlsruhe, Germany, #9090.3)
4% SDS (Carl Roth GmbH + Co. KG, Karlsruhe, Germany, #1057.1)
20% Glycerol (Carl Roth GmbH + Co. KG, Karlsruhe, Germany, #3783.1)
0,2% Bromophenolblue (Carl Roth GmbH + Co. KG, Karlsruhe, Germany, #A512.1)

### 2.2 Experimental animals

All animals used in this study were housed under constant temperature on a 12h light/dark cycle and had access to food and water *ad libitum* and were on a C57/BL6-J background. All mice were handled in strict adherence to local governmental and institutional animal care regulations.

### 2.3 Lysate Preparation

Cell lysates were prepared by direct lysis of cells in 2x SDS LB on the plate using a cell culture scraper and subsequent pulse-vortexing for 10 seconds. Sciatic nerves of either the right or the left side from two different mice were pooled and snap frozen in liquid nitrogen immediately after isolation. The harvested brain was divided into two pieces and lysates were prepared. Spleens were homogenized and shown data represent organs from four different mice (animal numbers 379, 383, 384 and 387). Tissues were homogenized using ceramic beads in a Precellys^®^ 24 homogenizer (Bertin Instruments, Montigny-le-Bretonneux,France) in either Pierce RIPA buffer (Thermo Fisher Scientific Inc., Waltham, MA, USA) with cOmplete protease inhibitor and phosSTOP phosphatase inhibitor (Roche Diagnostics GmbH, Mannheim, Germany) or in 2x SDS LB.

### 2.4 BCA Assay

The microscale BCA assay (Micro BCA Protein assay kit (Thermo Fisher Scientific Inc., Waltham, MA, USA, #23225) was performed according to the manufacturers instructions and a linear equation based on the linear trendline of the standard curvewas generated with Microsoft Excel and used for the determination of protein concentrations.

### 2.5 Dot blot and Ponceau S staining

Protein lysates or purified BSA were applied point wise to dry nitrocellulose membranes. Lysates in 2x SDS LB were boiled for 8 minutes at 98°C before applying them to the membrane. The lysates were allowed to dry on the membrane for 15 minutes and were either directly used for Ponceau S staining or concerning the samples in 2x SDS LB, the membranes were washed 3x 5min in deionized (DI)water on a shaker. Afterwards Ponceau S solution (0.1% Ponceau S in 5% acetic acid) was applied for one minute on loaded membranes and equal distribution was ensured. Afterwards the membrane was briefly washed with DI water until background staining was removed and membranes were placed into a plastic foil and scanned with a Epson Perfection V750 Pro scanner using the professional mode and the reflective document type in the scanning software.

### 2.6 Protein quantification with Fiji and Microsoft Excel

After creating a greyscale 8 bit image in the free, open-source Fiji software, the rectangle tool and ROI manager were used to define the different dots as regions of interest. The rectangle was always left at the same size for all dots to avoid variation in the “area” variable of the formula for the integrated density. After selection of all dots the pre-selected integrated density was measured and used for quantifications. Values were averaged from technical dupli-or triplicates and divided by 10^5^ for easy handling. A standard curve was generated using a linear “scatter chart” in Microsoft Excel and a linear trendline was inserted. The corresponding linear equation was used for the calculation of protein concentrations.

### 2.7 Cost calculations

We calculated the costs regarding each the PDB-method and the micro BCA assay for a reaction with 12 biological samples and standards. Both the commercial BCA assay and the selfmade variant (Reagent A: 1% sodium bicinchoninate, 2% sodium carbonate, 0.16% sodium tartrate, 0.4% NaOH, 0.95% sodium bicarbonate, 10M NaOH, pH 11.25, Reagent B: 4% cupric sulfate) were taken into the comparison. We provide a range of possible total costs, which depends on the distributor providing the ingredients.

We calculated the costs for the RIPA buffer and the SDS Gel loading buffer per 1ml. We compared commercial RIPA buffer (RIPA Lysis and Extraction Buffer, #89900, Thermo Fisher Scientific Inc., Waltham, MA, USA) and selfmade RIPA buffer (25mM Tris-HCl, pH 7.6, 150mM NaCl, 1% NP-40, 1% sodium deoxycholate, 0.1% SDS) with selfmade SDS Gel loading buffer (100 mM Tris-HCl, 4% SDS, 0.2% bromophenol blue, 20% glycerol). We generally chose high quality distributors, and the smallest available packing size of each product served as the basis for our calculations. Everyday lab chemicals such as Tris-HCl or NaCl were not included into the calculations. Included in the final costs for both commercial and selfmade RIPA buffer were the relative costs for a protease (c0mplete Protease Inhibitor, CO-RO Roche, Merck KGaA, Darmstadt, Germany) and phosphatase inhibitor (PhosStop, Phoss-Ro Roche, Merck KGaA, Darmstadt, Germany).

### 2.8 Immunoblotting

Immunoblotting was performed as previously described(7). The used antibodies are listed in Table 1. Blots were developed with Pierce ECL Western Blotting Substrate (Thermo Fisher Scientific Inc., Waltham, MA, USA)

**Table 1.** Cost estimations for the PDB assay as well as commercial and selfmade BCA kits.

| Method | Materials | Price (for 12 samples) |
| --- | --- | --- |
| Ponceau Dot Blot | • Ponceau powder<br>• Nitrocellulose membrane (84cm<sup>2</sup>)<br>• Bovine serum albumin | 2,05€ |
| Commercial BCA Kit | • Kit Materials<br>• Pierce 96-wellplate | 13,47€ |
| Selfmade BCA Kit | • Kit Materials<br>• Pierce 96-wellplate | Ranging from 15,29€ to 24,91€ (depending on producer) |
|  | Price for 1000 samples<br>Ponceau Dot Blot: 170,83€<br>Commercial BCA Kit:<br>~1122,50€<br><u>$\Delta$ = ~951,67€</u> |  |

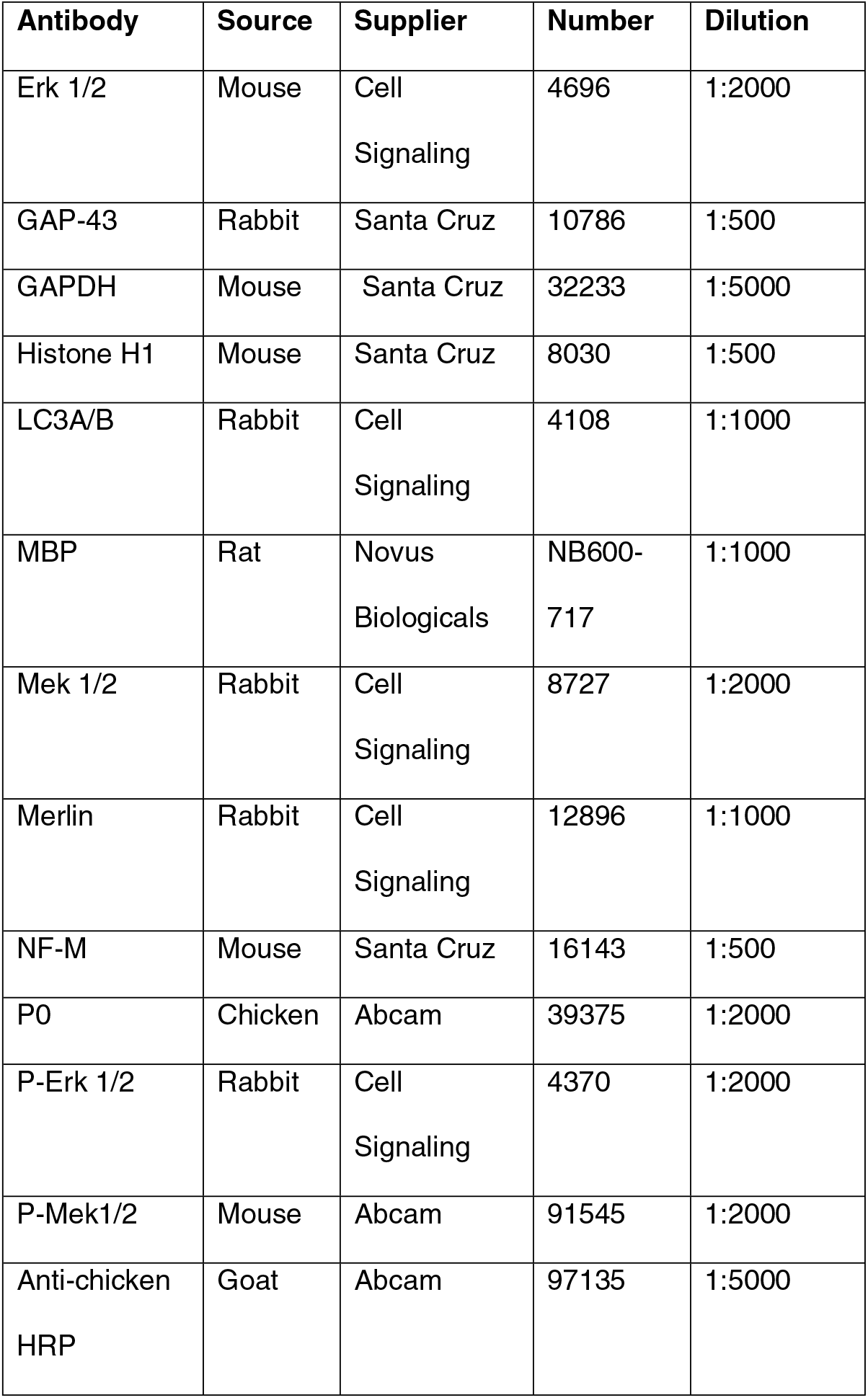
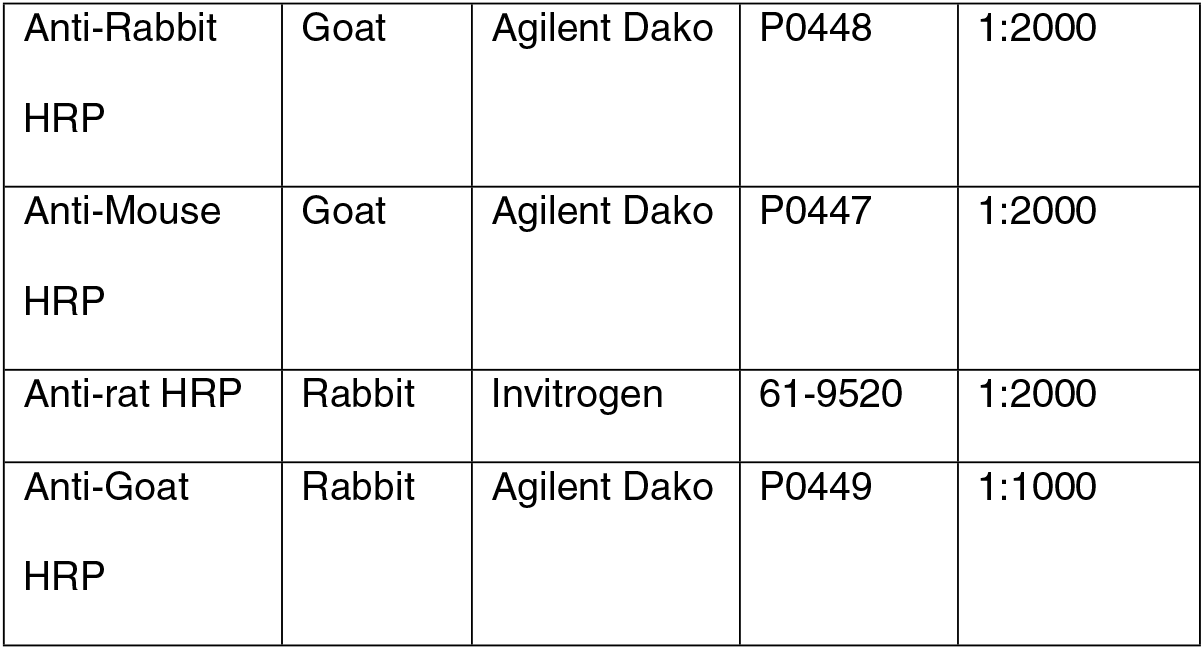

### 2.9 Statistical procedures and Figure preparation

Two-tailed, unpaired Student’s t-tests for calculation of p-values and F-tests to check the normal distribution of datasets were performed using Graphpad Prism 7.0. Statistical significance was accepted at p≤0.05. All data are presented as mean +/− SD. All figures were either made with Graphpad Prism 7.0 or Microsoft PowerPoint and were assembled in Adobe Photoshop CS6.

## 3. Results and Discussion

To test whether Ponceau Staining of Dot blots can be used as a protein quantification method we spotted undiluted, commercially available BSA solution at a concentration of 2 mg/ml in a range of 0.25 μg to 4 μg to nitrocellulose membranes. Membranes were stained with a 0.1% Ponceau S solution in 5% acetic acid for one minute and scanned to quantify the stained dots (**Fig. 1A**).

**Figure 1.**
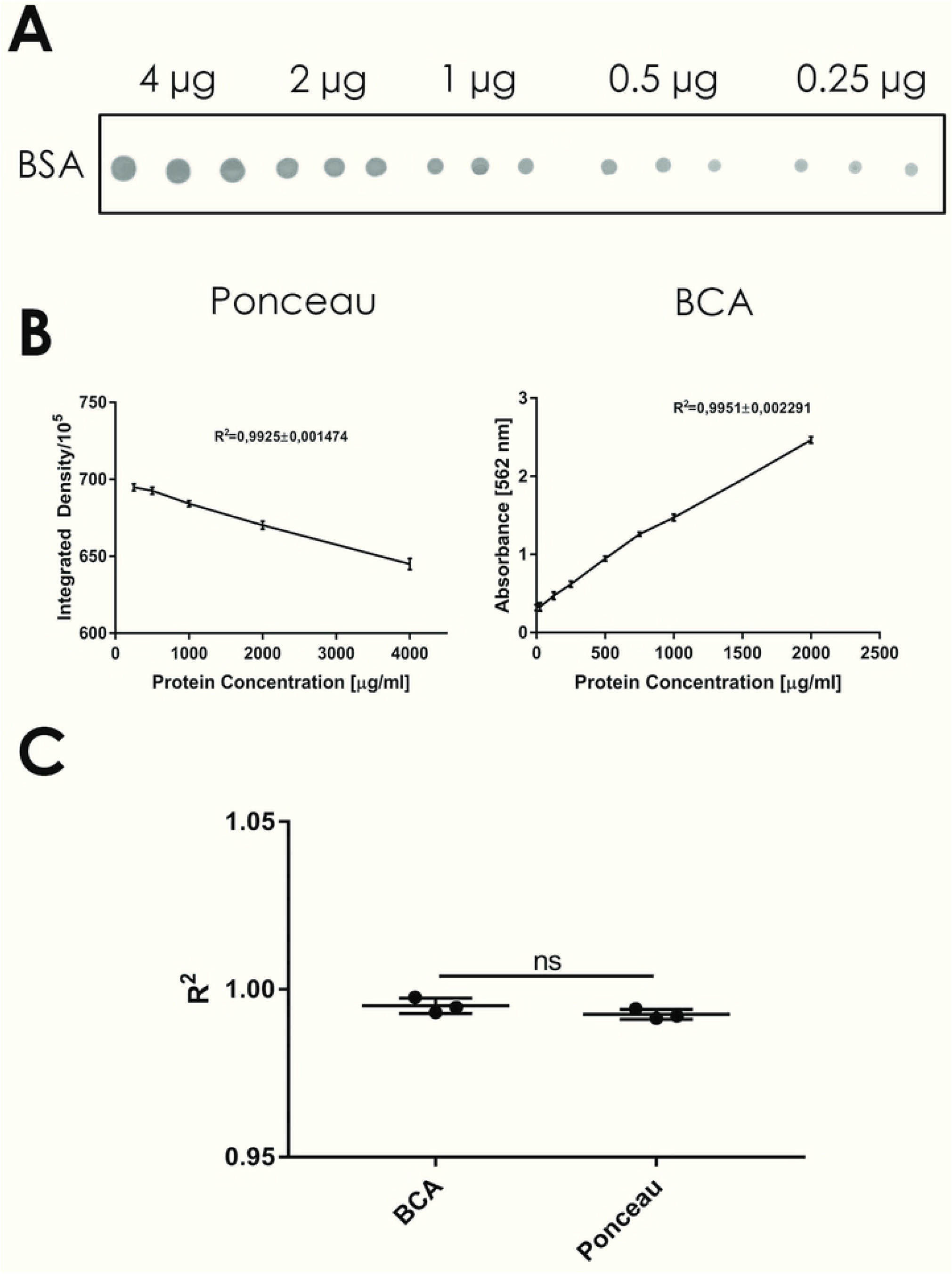
**A** Representative Ponceau S stained dot blot of undiluted BSA spots of indicated protein amounts (n=1). Different amounts of BSA were applied onto the membrane in triplicates. **B** Linear standard curves of different Ponceau S stained dot blots from BSA standards generated with either PDB assay or BCA assay (n=3, each). Replicates are defined by usage of BSA from three different ampules. **C** Comparison of correlation coefficients from either BCA or PDB assay (n=3, ns=p>0.05, unpaired, two-sided student’s t-test).

Because the sizes of the resulting dots were unequal we used the “Integrated Density” of each dot which is the product of the selected area and its mean grey value over the measured area. By using the same rectangle size when analysing the dots in *Fiji* only the grey values decrease as the protein amount increases in a linear manner (**Fig. 1B, left**). The resulting linear standard curve revealed high consistency/low variability within the three different standards. A correlation coefficient *R^2^* of 0,9925 also indicated high linearity of the standard curve generated with our PDB-assay. To avoid the necessity of spotting different volumes, which could be a reason of variation, we diluted the BSA in either ddH2O or the same RIPA buffer in which we lysed the different organs used in this study. Dilution of the BSA in ddH_2_O still gives a clear dot, but unfortunately the dilution of BSA in RIPA buffer resulted in the distribution of the BSA in form of circles(**Fig.S1**).

**Figure S1**

Ponceau S stained dot blot of BSA diluted either in ddH_2_O or RIPA buffer. Diluted samples were applied onto the membrane in duplicates.

This is known in the literature as “coffee rings” (8). This could not be prevented by washing the membrane after spotting the BSA onto it, to remove potentially interfering SDS. It has been shown that the formation of such a “coffee rings” depends on the evaporation speed of the liquid and the particle movement. We can only speculate about the cause of this observation but we assume that the low concentration of SDS inside the RIPA buffer decreases the speed of the particle movement, as it has been shown that lower amounts of SDS decrease the diffusion coefficient of ovalbumin (9).The dilution of BSA in ddH_2_O also resulted in a non-linear standard curve, making it inappropriate or unsuitable for proper quantification of lysates (**Fig. S2**).

**Figure S2**

**A** Representative Ponceau S stained dot blot of in ddH_2_O diluted BSA of indicated amounts (n=1). Different amounts of BSA were applied onto the membrane in triplicates.

**B** Linear standard curve of different BSA standards diluted 1:1 in ddH_2_O (n=3).

This is probably the result of loss of diluted BSA during serial dilution of the standard on the walls of pipette tips and tubes (10). Therefore, we used undiluted BSA to generate standard curves. For comparison, we also generated the standard curve by bicinchoninic acid (BCA) assay, a well established method. The BCA assay also showed a good linearity within a range of 125 ng to 2 μg.(**Fig. 1B, right**). Comparison of the correlation coefficients demonstrates that the PDB assay is in performance completely equal to a BCA assay (**Fig. 1C**).

However, the commonly used microscale BCA assay is known to display saturation of the photocolorimetric reaction at protein concentrations outside the range of the standard, meaning above 2 μg/μl.The PDB has the big advantage of being linear also in the range of higher amounts of protein (in this experiment up to 4 μg).

To test the applicability of our PDB method with tissue lysates, we collected spleens as protein-rich organs from four different mice, lysed them in 1 ml RIPA buffer and used 1 μl per dot of this lysate for quantification (**Fig. 2A**).

**Figure 2.**
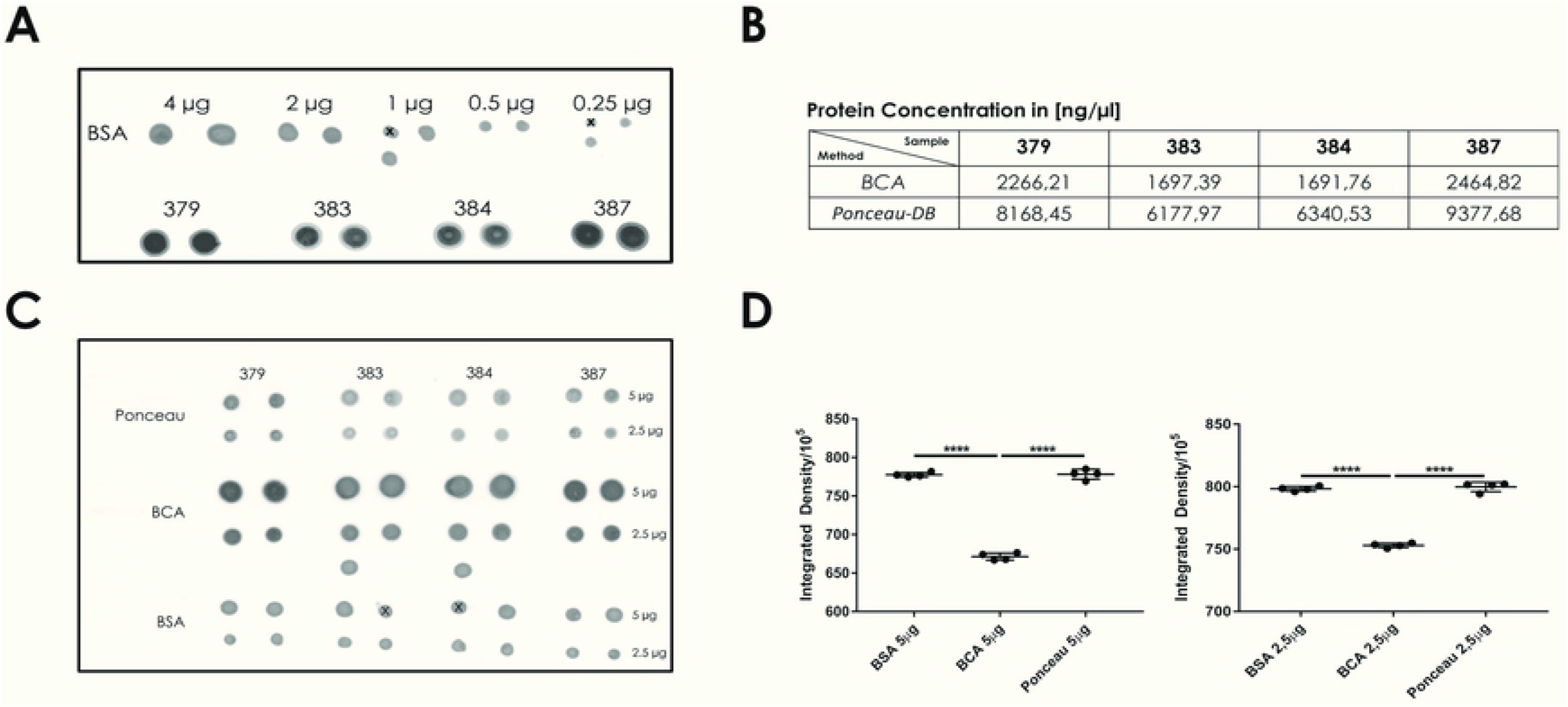
**A** Membrane with spotted dots of either BSA standard (range from 0,25μg to 4μg as indicated handwritten) and spleen lysates from four different mice (#379, #383. #384, #387). Different samples/amounts of protein were applied onto the membrane in duplicates. Dots crossed out with an ”x” were excluded due to accidental application of unequal BSA amounts. **B** Table displaying protein concentrations of spleen lysates determined by either PDB or BCA assay. **C** Membrane with 5μg or 2,5μg dots of spleen lysates or BSA (lower two rows). For the determination of the protein concentrations inside lysates the PDB assay was used for the dots in the first two rows and the BCA assay in row three and four. Different samples/amounts of protein were applied onto the membrane in duplicates. **D** Diagrams depicting integrated densities of dots from the membrane shown in **C** (n=4, ****=p<0.0001, two-sided, unpaired student’s t-test).

In parallel, the same lysates were quantified by BCA assay for comparison. Due to the high protein content in the spleen lysates, the BCA assay showed values around the upper border of the standard range, between 1,7 μg/μl and 2,4 μg/μl. In contrast, with the PDB assay the protein concentrations were determined between ~6 and 9 μg/ μl, three to four times higher than the values given by the BCA method (**Fig. 2B**). To validate the concentrations determined by PDB, we calculated the required lysate volume for 5 and 2,5 μg with the concentrations from BCA and PDB and applied these amounts together with 5 and 2,5 μg BSA onto membranes (**Fig. 2C**). Strikingly, the staining of dots of lysates which concentrations were determined with the BCA assay were much stronger than those which have been quantified by PDB. This is reflected in **Fig.2D:** Dots of BCA quantified lysates displayed much lower integrated densities meaning higher protein content which can be explained by saturation of the BCA assay in working ranges above 2 μg. Integrated densities of PDB quantified dots were equal to those from BSA, indicating reliable performance of our method.

Although widely used RIPA buffer has the disadvantage that around 10-30 % of all proteins are lost during lysis due to the insolubility of some proteins in RIPA buffer(11). Hence, we tested another commonly used lysis buffer, 2x SDS Gel loading buffer, containing 4% SDS for more efficient solubilization of test tissues (11). We first applied BSA, diluted 1:1 in 2x SDS LB, to a membrane and compared it to the staining of BSA diluted in ddH_2_O. A nearly invisible circular shape of the applied dot was observed which was in comparison to the strong signal of the same amount of BSA diluted in ddH_2_O nearly nothing (**Fig. S3**).

**Figure S3**

Ponceau S stained dot blot of BSA diluted either in ddH_2_O or 2x SDS LB which was not washed in DI-tap water before Ponceau S staining. Diluted samples were applied onto the membrane in duplicates.

We hypothesized that the decreased staining effectivity might be due to the high concentration (2%) of SDS in the 1:1 diluted sample which could interfere with the binding of the Ponceau S dye to proteins. Therefore, we washed the membrane after drying three times for five minutes in DI-tap water before staining. This led to effective staining of the dots which were, compared to dots of undiluted BSA, weaker in their intensities but spread over a larger area. This was probably due to the fact that two times the volume of undiluted BSA was used to achieve equal protein amounts (**Fig. 3A**). Interestingly, we did not again observe the “coffee ring”-phenomena as we did when we diluted the BSA in RIPA buffer. Again we can only speculate and explain this by the reported observation that higher amounts of SDS normalize the diffusion coefficient of ovalbumin which was decreased by low amounts of SDS(9). Therefore, the speed of evaporation of the lysate droplet is again equal to the speed of the particle movement within the droplet. As before, we used the same BSA standards as in **Fig.1** to produce a standard curve of BSA diluted in 2x SDS LB. This mean standard curve also again displayed good linearity, reflected by a mean *R^2^* of 0,9945 (**Fig. 3B**). Compared to the other correlation coefficients shown in **Fig. 1C**, there were no changes between all three different methods in linearity of prepared standards as adressed by the correlation coefficients (**Fig. 3C**).

**Figure 3.**
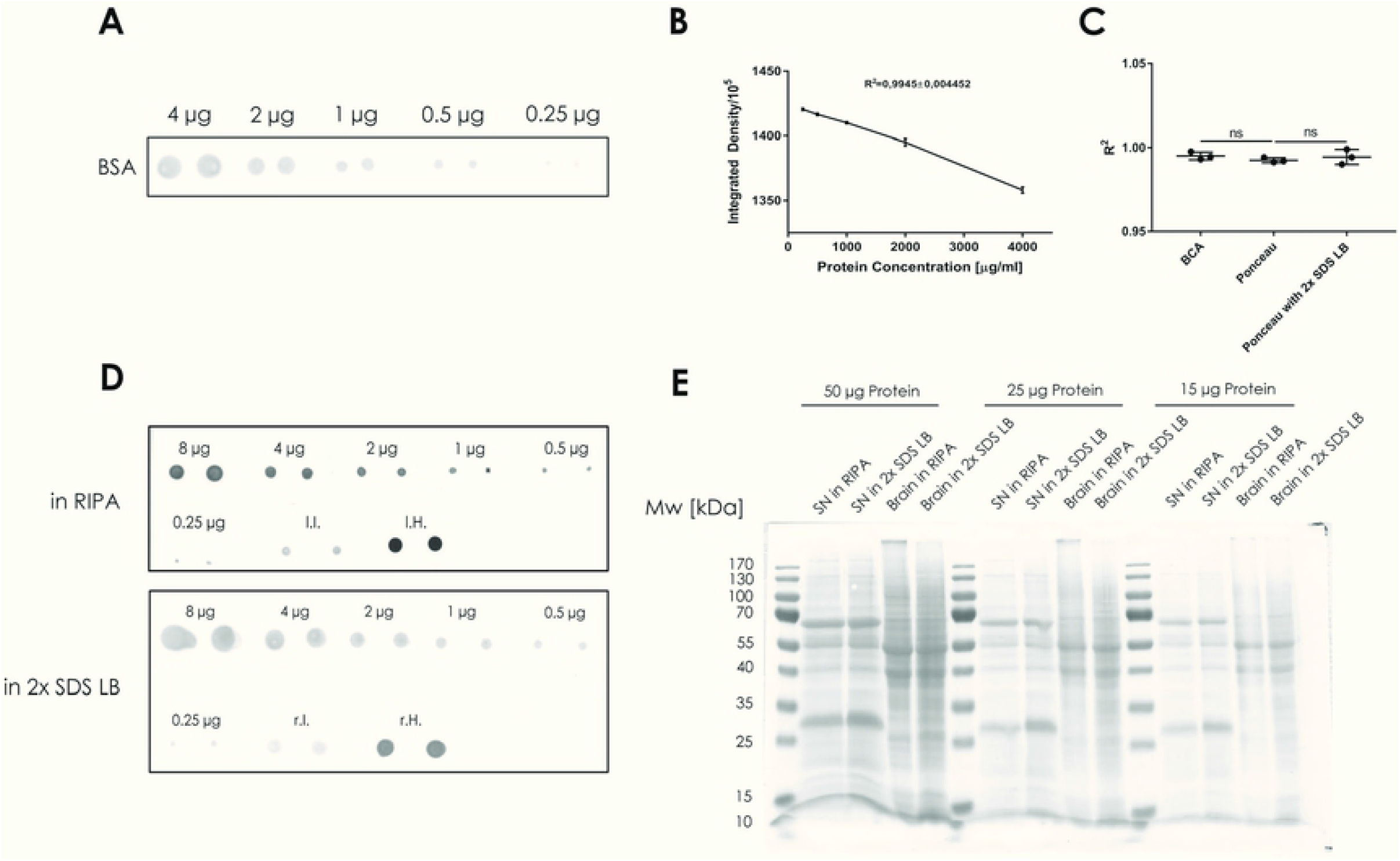
**A** Representative Ponceau S stained dot blot of in 2x SDS LB diluted BSA of indicated amounts (n=1). Different amounts of BSA were applied onto the membrane in duplicates. **B** Linear standard curve of different Ponceau S stained dot blots from BSA standards diluted 1:1 in 2x SDS LB (n=3). Replicates are defined by usage of BSA from three different ampules. **C** Comparison of correlation coefficients from standard curves of BCA assay, PDB assay with undiluted BSA or PDB assay with BSA diluted 1:1 in 2x SDS LB (n=3, ns=p>0.05, unpaired, two-sided student’s t-test). Please note that the data for the BCA assay and PDB assay with undiluted BSA are the same as shown in **Fig.1C**. **D** Membranes with stained undiluted BSA standard curves (ranging from 8μg to 0,25 μg), 1μl spots of sciatic nerve (“I.I.”), brain (“l.H.”) lysates in RIPA buffer (upper membrane) or in 2x SDS LB 1:1 diluted BSA (ranging from 8μg to 0,25 μg), 2μl spots of sciatic nerve (“r.I.”) and brain (“r.H.”) lysates in 2x SDS LB buffer diluted 1:1 in ddH_2_O (lower membrane). Different samples/amounts of protein were applied onto the membrane in duplicates. **E** Ponceau S stained membrane with different protein amounts of sciatic nerve lysates in either RIPA buffer or 2x SDS LB.

To test the suitability of direct tissue lysis in 2x SDS LB and to compare the extraction ability with RIPA, we used sciatic nerves and a brain from C57/BL6 mice. These tissues are normally hard to lyse due to their high content of fatty myelin. We pooled the sciatic nerves from the left and right side of two different mice (left sciatic nerves were lysed in RIPA buffer and right sciatic nerves were lysed in 2x SDS LB) and also used one mechanically disrupted mouse brain which we divided into two halves and subsequently lysed either in RIPA buffer or 2x SDS LB. Since we expected very high concentrations for the lysed brain, we also included 8 μg of BSA into the range of our standard. The resulting standard curve still maintained a good linearity (**Fig. S4**), supporting the suitability of Ponceau S dye for the quantification of tissue lysates with high protein content.

**Figure S4**

Linear standard curves of Ponceau S stained dot blots from BSA standard shown in **Fig.3D** (n=1).

With both buffers we could lyse and determine protein concentrations effectively of brain pieces and nerves (**Fig. 3D**). To test whether quantification of protein concentrations inside lysates, in which proteins were differentially extracted, gave us in both cases true values, we subjected the lysates of 50, 25 and 15 ug protein content to a SDS-PAGE followed by protein transfer to the membrane, subsequent Ponceau S staining and immunoblotting.

The Ponceau S staining showed overall equal loading between the two extraction methods if one compares only sciatic nerves or brain lysates among themselves (**Fig. 3E**). This proves similar performance of the PDB assay with either lysates prepared in in RIPA buffer or in 2x SDS LB (**Fig. 3E**). Intriguingly, there was a general difference between the loading of sciatic nerves and brain lysates (**Fig. 3E**). We suppose that this is due to the high abundance of albumin (strong band below 70 kDa) and IgG heavy (strong band slightly above 55 kDa) and light (strong band between 25 and 35 kDa) chain in the PNS which are absent in the CNS due to the blood brain barrier (12, 13). The presence of these highly abundant serum proteins would lead to overestimation of the real protein content of the sciatic nerve itself and therefore leads to unequal loading compared to both brain lysates. This is an important point if researchers attempt to compare expression of different proteins between CNS and PNS.

As expected, subsequent immunoblotting revealed better extraction of different proteins by 2x SDS LB. It has been described that e.g. cytoskeleton associated proteins and extracellular matrix components are to a certain degree insoluble in RIPA buffer(11). The tumor suppressor protein merlin as a cytosekeleton associated protein was extracted more in 2x SDS LB in both sciatic nerve and brain lysates as described before (14)(**Fig.4**). Cytoplasmic proteins like MEK 1/2, ERK 1/2, GAPDH and GAP-43 were present to the same extent in both lysates. The nuclear protein Histone H1 and the autophagic vesicle membrane proteins LC3A/B were slightly less abundant in the RIPA buffer extractions. Since we included phosphatase inhibitors in the RIPA buffer, we were surprised as we detected slightly higher P-ERK1/2 but massively higher P-MEK1/2 signals suggesting more efficient phosphatase inhibition in 2x SDS LB, probably due to the strong denaturing effect of SDS. This finding is important for researchers studying fast-changing signaling processes e.g. during nervous system regeneration and degeneration (15, 16). Lastly, we observed a slightly enhanced ability of 2x SDS LB to extract the axonal intermediate filament neurofilament-M and obviously enhanced ability to solubilise the extracellular matrix associated myelin proteins myelin basic protein (MBP), which is present in both the PNS and CNS, and the PNS specific myelin protein zero (P0; **Fig. 4**).

**Figure 4.**
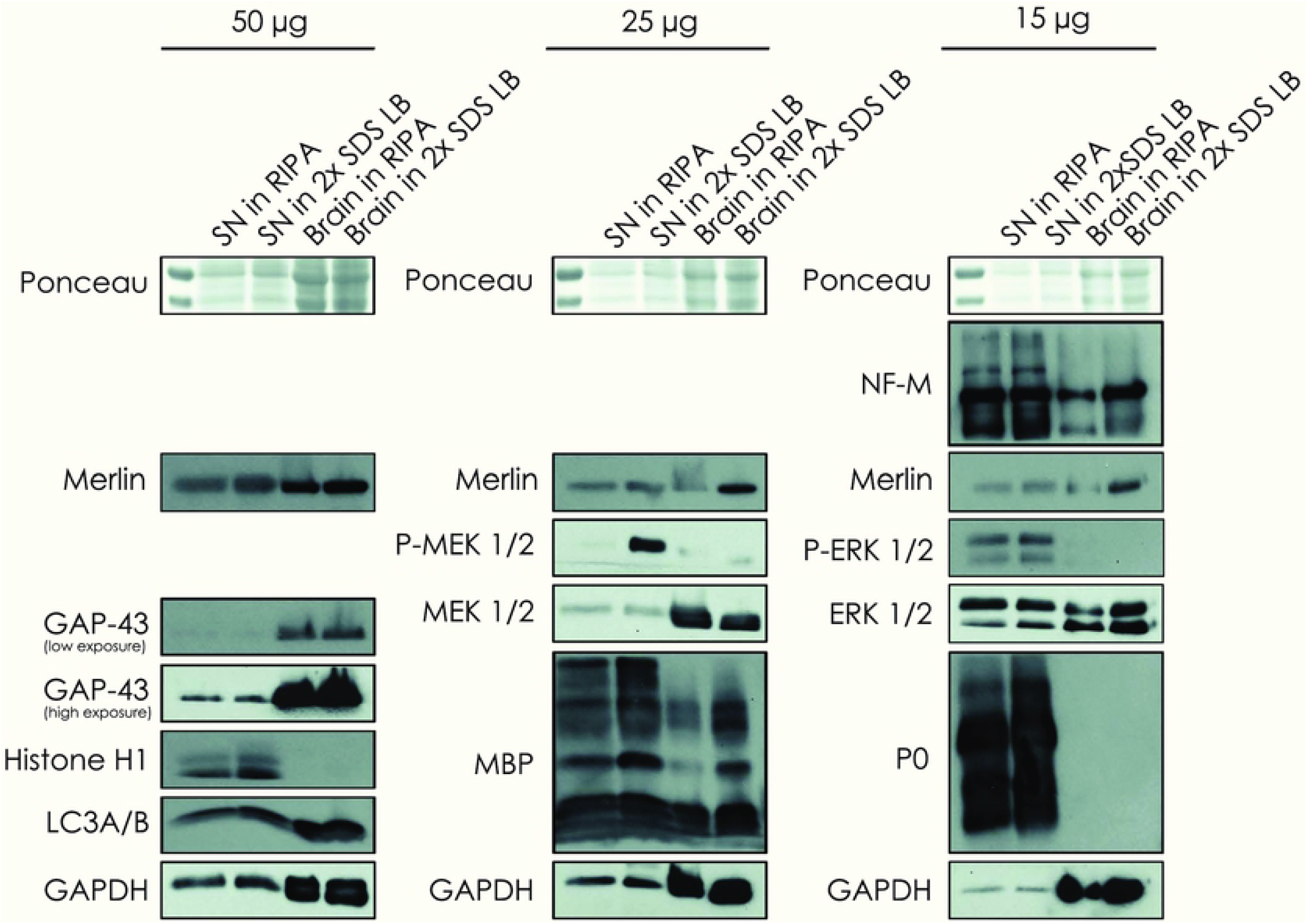
Immunoblots for indicated target proteins of the membrane shown in **Fig.3E** (n=1).

Throughout the course of our experiments we realized that the PDB assay, especially if combined with the use of 2x SDS LB as lysis buffer instead of RIPA buffer, must be relatively cheap if compared to the established workflow in our laboratory from tissue harvesting to immunoblotting. Therefore, we estimated the possible amount of money a laboratory could save with the usage of our method. First we calculated the costs for one reaction with quantification of 12 samples. While a “selfmade” BCA kit would cost between 15,29€ and 24,91€ and a commercial BCA kit 13,47€ our PDB assay only costs 2,05€ per reaction. Based on our laboratory experience we know that a prepared Ponceau S solution can be used at least 20 times to stain membranes.One bottle with 10g of Ponceau S powder at a final dilution of 0,1% in 5% acetic acid is enough to stain 800 membranes with 12 samples per membrane. If we calculate the costs for measuring 1000 samples and compare these with the costs of 1000 BCA reactions, it turns out that with our PDB method a laboratory would save around 951,67€ (**Tab.1**).

Since we have shown that direct lysis of tissues in 2x SDS LB is not only compatible with our method but rather even more recommended because it extracts and solubilises different proteins better (**Fig.4**), as it has been reported previously(11), we also calculated how much money could be saved with the usage of 2x SDS LB instead of RIPA buffer with addition of phosphatase and protease inhibitors. One ml RIPA lysis buffer with phosphatase and protease inhibitors routinely used in our laboratory costs around 2,41€ while the same volume of selfmade 2x SDS LB only costs 0,1€. One ml of selfmade RIPA lysis buffer with phosphatase and protease inhibitors would still cost 1,48€.. If we project this to 1000 prepared lysates, a laboratory would save between 1380 and 2310€ (**Tab.2**). Adding up these amounts shows that the laboratory could save 2331,67 and 3261,67€ per 1000 lysates by using 2x SDS LB for lysis of tissues and the PDB assay for quantification of these lysates (**Tab.2**).

**Table 2.**
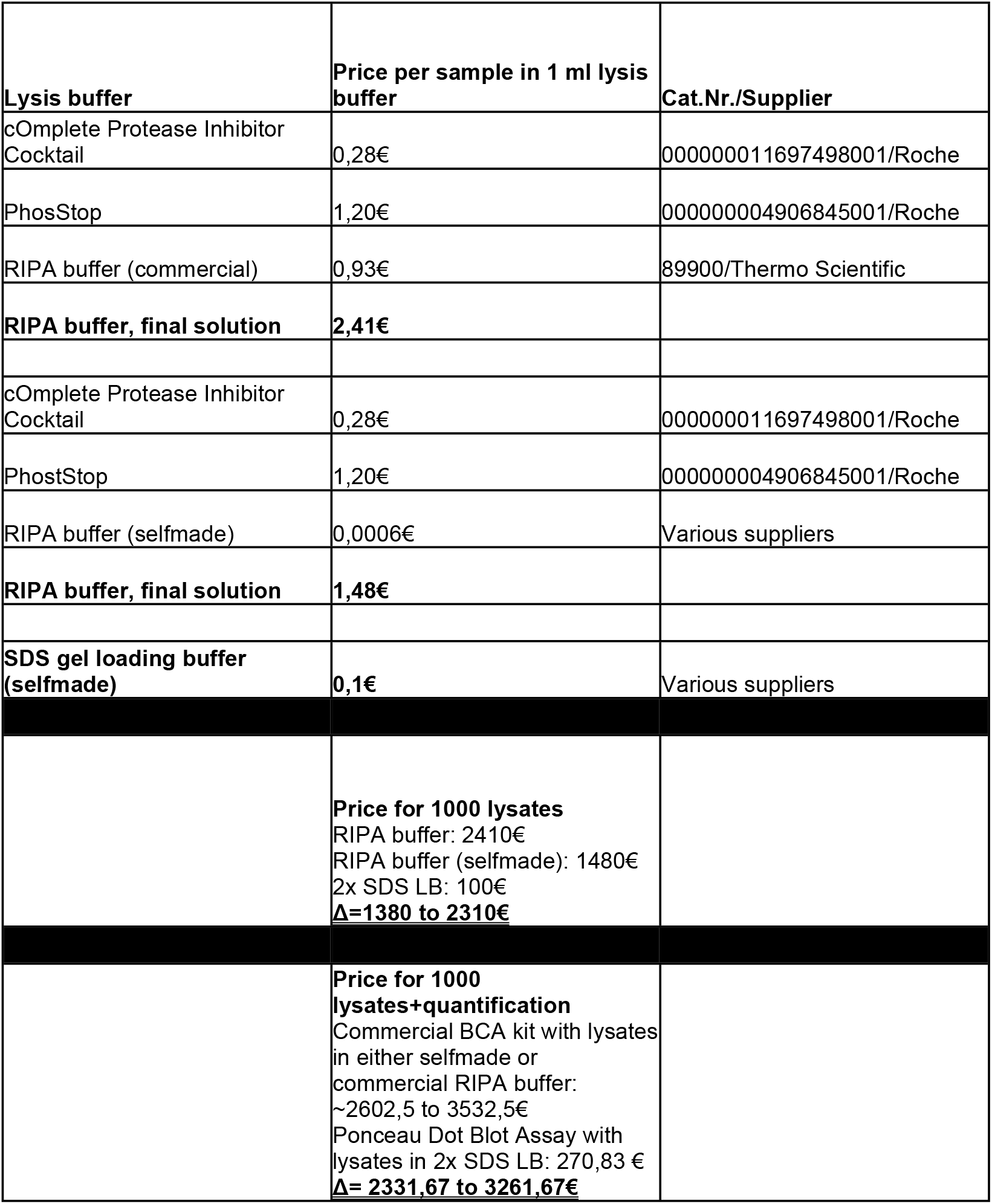
Cost estimations for different lysis buffers used with either our PDB assay or a BCA assay.

Although a similar principle was described previously (17), our study highlights some more critical points and adds a new improvement: If using RIPA buffer as lysis buffer it is extremely important not to dilute the BSA which is used for preparation of a standard curve. This also leads to a faster workflow of our method compared to the other one reported. If BSA is diluted in ddH_2_O, a non-linear standard curve will be the result and if diluted in RIPA buffer, a circle instead of a dot will form. Furthermore, we could show that tissues can directly be lysed in 2x SDS Gel loading buffer which is faster and superior to lysis in RIPA buffer and the preferable method since loosing a lot of protein during extraction and lysis could be avoided. The fact that Ponceau S staining of dot blots is suitable for accurate quantification of tissue lysates is another important improvement compared to the publication from Morcol et al., who only used different purified proteins but no tissue lysates(17).

## 4. Conclusion

We describe a rapid, low-budget and highly reliable technique for quantification of protein lysates as an alternative to more common established methods like the BCA assay. Our method is a considerable improvement of the method described previously (17), based on the aforementioned points.

Different protein extractions and lysis protocols could also be the reason for contradicting reports in the literature. Since Western blotting with subsequent immunodetection of different target proteins is one of the most widespread methods in biomedical research, laboratories working on the same model system/organ or topic of interest could standardize the obtainment of results by using the same strategies/protocols which would lead to the publication of more reliable results. With the money saved by the usage of our technique, the research in every laboratory could be highly improved by the contingency to purchase more antibodies, chemicals, biological materials, etc.

## Acknowledgements

The authors would like to thank Debra Weih for critical reading and editing of the manuscript.

